# Cdk8 directly regulates glycolysis via phosphorylation of Gcr2

**DOI:** 10.1101/2021.03.12.435012

**Authors:** Maria J. Aristizabal, Eoghan O’Duibhir, Wim de Jonge, Kristy Dever, Nicole Hawe, Marian Groot Koerkamp, Mariel Brok, Dik van Leenen, Philip Lijnzaad, Michael S. Kobor, Frank C.P. Holstege, Ivan Sadowski

## Abstract

*CDK8* encodes an evolutionarily conserved Mediator complex kinase subunit that functions in general and context-specific transcription regulation by phosphorylating core components of the transcription machinery and gene-specific transcription factors. To better understand the role Cdk8 in transcription regulation, we performed high-resolution gene expression time course analysis following nuclear depletion of Cdk8. Focusing on the earliest gene expression alterations revealed dysregulation of genes encoding glycolysis enzymes, suggesting a functional link to Gcr1 and Gcr2, key transcriptional activators of these genes. Consistently, we found that nuclear depletion of Cdk8 altered the mRNA levels of glycolysis genes as well as the promoter occupancy of Gcr2, but not Gcr1. Examination of the Gcr2 protein sequence revealed a putative Cdk8 phosphorylation site at serine 365, which we confirmed using *in vitro* and *in vivo* assays. Importantly, phospho-mutant *GCR2* recapitulated the growth and gene expression defects of the *GCR2* deletion mutant, effects not observed with a phospho mimetic mutant. As such, our work highlights Gcr2 as a new Cdk8 substrate, revealing that its phosphorylation is critical for the activation of genes encoding glycolysis enzymes.

## INTRODUCTION

Cdk8 is a cyclin dependant kinase that together with Cyclin C, Med12 and Med13 forms the kinase submodule of the RNA Polymerase II-associated Mediator complex. Mediator is a conserved transcription co-activator complex required for the response to transcription factors and is likely involved in the transcription of all protein-coding genes (reviewed in (Conaway and Conaway, 2011; Kornberg, 2005).

Despite extensive research, the role of Cdk8 in transcriptional regulation is still enigmatic. Evidence indicates that Cdk8 has both general and gene-specific roles in transcription, and can act as a positive or negative regulator of gene expression in both yeast and metazoan cells. Observations demonstrating an association with the Mediator complex, and genetic links to the carboxy-terminal domain (CTD) of RNA polymerase II (Aristizabal et al., 2013; Kim et al., 1994; Liao et al., 1995), position Cdk8 as a general modulator of transcription. Accordingly, Cdk8 phosphorylates components of the general transcription machinery including the RNA polymerase II CTD (Liao et al., 1995), other Mediator subunits (Hallberg et al., 2004; Liu et al., 2004; Miller et al., 2012; Peppel et al., 2005; Poss et al., 2016), as well as the TATA Binding / TFIID-associated proteins Taf2 and Bdf1 (Liu et al., 2004).

Several lines of evidence also support gene-specific activities for Cdk8. First, in stark contrast to most subunits that make up the Mediator core, genes encoding subunits of the kinase submodule are dispensable for viability in yeast (Hengartner et al., 1995). In fact, mutant alleles of *cdk8*, cyclin C, *med12* or *med13* have been identified in over 10 different genetic screens aimed at identifying modulators of specific regulatory processes (Balciunas and Ronne, 1995; Carlson et al., 1984; Li et al., 2005; Surosky et al., 1994; Tabtiang and Herskowitz, 1998). Second, high-throughput gene expression profiling revealed that only a subset of genes were affected upon *cdk8* deletion, compared to genome-wide effects observed upon disruption of Mediator core components like *MED17* (Holstege et al., 1998; Peppel et al., 2005). Third, multiple gene-specific transcription factors are direct targets of Cdk8 (Chi, 2001; Hirst et al., 1999; Lenssen et al., 2007; Nelson et al., 2003; Raithatha et al., 2011). Amongst these, Gcn4, Ste12 and Phd1 are negatively regulated by Cdk8-dependent phosphorylation, which promotes their proteolytic degradation (Chi, 2001; Nelson et al., 2003; Raithatha et al., 2011; Rosonina et al., 2012). In contrast, Cdk8-dependent phosphorylation of the Gal4 transactivator is required for full induction of *GAL* gene expression, underscoring both negative and positive effects of Cdk8 on specific gene transcription (Hirst et al., 1999).

Most of what we know about Cdk8 has emerged from studies in yeast using steady-state perturbation *via* gene deletion. Knockout of *CDK8* in yeast has at least two obvious phenotypic consequences under standard growth conditions, flocculence (Holstege et al., 1998) and reduced growth rate (Hengartner et al., 1995). Flocculence is produced by a differentiated state, known as filamentous or pseudohyphal growth, wherein yeast form elongated cells that remain attached after cell division due to changes in the composition of their cell wall in response to nutrient limitation (reviewed in (Cullen and Sprague, 2012). Accordingly, Cdk8 is known to regulate filamentous growth by phosphorylating the transcription factors Ste12 and Phd1, modulating their stability (Nelson et al., 2003; Raithatha et al., 2011). Slow growth, has the potential to confound the effect of *CDK8* deletion on gene expression, an effect that has been observed for other null mutants (O’Duibhir et al., 2014). Thus, assessing a direct role for Cdk8 in gene regulation is complicated by a challenge common to many studies aimed at illuminating molecular mechanisms of transcription regulation, the potential for confounding secondary effects produced by null mutations. These can emerge from downstream events, or exposure of compensatory mechanisms that alter gene expression or result in a particular phenotypic state, including slow growth.

Reduced proliferation of *CDK8* null mutants in standard growth media containing glucose indicates that *CDK8*, although not essential, is required for the most rapid growth of yeast. While these effects likely emerge from defects in specific gene expression programs, they may confound gene expression profiles and mask direct effects of *CDK8* in transcription. To separate the effects of slow growth from direct effects of *CDK8* function we made use of rapid, non-invasive conditional nuclear depletion of Cdk8 using the “anchor-away” system (Haruki et al., 2008). Examining gene expression changes in high temporal resolution, immediately following Cdk8 nuclear depletion, revealed decreased expression of specific gene subsets, some of which had not been previously associated with Cdk8 function. Analysis of the regulatory regions of these genes pointed to Gcr1 and Gcr2, key transcriptional activators of genes encoding glycolysis enzymes (Uemura and Jigami, 1992a), as potential targets of Cdk8. Examining this relationship further revealed that Cdk8 directly phosphorylated the transcription factor Gcr2, a modification required for growth on glucose containing media and the expression of genes encoding enzymes involved in glycolysis.

## MATERIALS AND METHODS

### Yeast strains and plasmids

All yeast strains and plasmids used in this study are listed in Table 1 and Table 2. The parental anchor-away strain in the S288C/BY4742 background was described previously (de Jonge et al., 2017).

**Table 1.** Strains used in this study

| Strain | Genotype | Background | Source |
| --- | --- | --- | --- |
| MA-Y9 | Matalpha GCR2::KAN ade2-1 can1-100 his3-11 leu2-3,112 trp1-1 ura3-1 LYS2 pRS314[] | W303 | This study |
| MA-Y10 | Matalpha GCR2::KAN ade2-1 can1-100 his3-11 leu2-3,112 trp1-1 ura3-1 LYS2 pRS314[GCR2] | W303 | This study |
| MA-Y11 | Matalpha GCR2::KAN ade2-1 can1-100 his3-11 leu2-3,112 trp1-1 ura3-1 LYS2 pRS314[gcr2-S365A] | W303 | This study |
| MA-Y12 | Matalpha GCR2::KAN ade2-1 can1-100 his3-11 leu2-3,112 trp1-1 ura3-1 LYS2 pRS314[gcr2-S365D] | W303 | This study |
| BY-Parental-AA | tor1-1; Dfpr1; RPL13A-2xFKBP12-NATMX6; MET15; his3-1; leu2; lys2; ura3; MATalpha | BY4742 | (de Jonge et al., 2017) |
| BY-cdk8AA | tor1-1; Dfpr1; RPL13A-2xFKBP12-NATMX6; MET15; his3-1; leu2; lys2; ura3; cdk8-FRB-yEGFP-hnhMX6 MATalpha | BY4742 | This study |
| BY-cdk8AA GCR1-3V5-his | tor1-1; Dfpr1; RPL13A-2xFKBP12-NATMX6; MET15; his3-1; leu2; lys2; ura3; gcr1-3V5-HIS; cdk8-FRB-yEGFP-hnhMX6 MATalpha | BY4742 | This study |
| BY-cdk8AA GCR2-3V5-his | tor1-1; Dfpr1; RPL13A-2xFKBP12-NATMX6; MET15; his3-1; leu2; lys2; ura3; gcr2-3V5-HIS; cdk8-FRB-yEGFP-hnhMX6 MATalpha | BY4742 | This study |

**Table 2.** Plasmids used in this study

| Plasmid | Relevant Genotype | Source or Reference |
| --- | --- | --- |
| pMA6 | pRS314 | This study |
| pMA81 | pRS314[GCR2] | This study |
| pMA82 | pRS314[gcr2-S365A] | This study |
| pMA83 | pRS314[gcr2-S365D] | This study |
| pIS484 | *URA3 ARS-CEN TEF1p-Σ1278b CDK8-3XFLAG-His<sub>6</sub>* | (Nelson et al., 2003) |
| pIS529 | *URA3 ARS-CEN TEF1p-Σ1278b CDK8D290A-3XFLAG-His<sub>6</sub>* | (Nelson et al., 2003) |

### Cell growth

Strains were streaked from −80°C stocks onto appropriate selection plates and grown for 3 days. Liquid cultures were inoculated from independent colonies and grown overnight in Synthetic Complete (SC) medium: 2gr/L Drop out mix Complete and 6.71gr/L Yeast Nitrogen Base without AA, Carbohydrate & w/AS (YNB) from US Biologicals (Swampscott, USA) with 2% D-glucose. Two cultures each of both parental anchor-away (AA) and Cdk8-AA strains were grown from overnight cultures. For growth rate determination, cells were grown in 1.5 ml of SC media at 30°C with shaking at 230 rpm in an automated Infinite 200 incubator (Tecan). For time course experiments and protein isolation, overnight cultures were diluted to an OD_600_ of 0.15 in 50 ml of fresh medium and grown in 250 ml Erlenmeyer flasks at 30°C with 230 rpm in a shaking incubator to an optical density of 0.6. Rapamycin was added to each of the four cultures giving a final concentration of 7.5μM (time point 0). Cultures were sampled for mRNA, flocculence and OD_600_ measurements at the indicated time points. Throughout the time course, cultures were diluted with fresh SC media containing 7.5μM rapamycin in order to maintain an OD_600_ of 0.4 to 1.4, thus avoiding entry into the diauxic shift.

### RNA extraction

Cell were harvested by centrifugation (4000 rpm, 3 min) and pellets were immediately frozen in liquid nitrogen. Total RNA was prepared by phenol extraction and cleaned using a customized Sciclone ALH 3000 Workstation. Briefly, frozen cells (−80°C) were resuspended in 500 μl Acid Phenol Chloroform (Sigma, 5:1, pH 4.7), followed by immediate addition of an equal volume of TES-buffer (TES: 10 mM Tris pH 7.5, 10 mM EDTA, 0.5% SDS). Samples were vortexed vigorously for 20 seconds, incubated in a water bath for 10 minutes at 65°C and vortexed vigorously one more time. Samples were then placed in a thermomixer (65°C, 1400 rpm) for 50 minutes followed by 20 min of centrifugation (4°C, 14000 rpm). Phenol extraction was repeated once, followed by a Chloroform:Isoamyl-alcohol (25:1) extraction. RNA was precipitated with Sodium Acetate (NaAc 3M, pH 5.2) and ethanol (96%, −20°C). The RNA pellet was washed with Ethanol, dissolved in sterile water (MQ), snapfrozen and stored at −80°C. All strains were profiled four times from two independently inoculated cultures.

### Microarray profiling

Dual-channel 70-mer oligonucleotide arrays were employed with a common reference wild type RNA. All steps after RNA isolation were automated using robotic liquid handlers. These procedures were first optimized for accuracy (correct fold change (FC)) and precision (reproducible result), using a spiked-in RNA calibration (Bakel and Holstege, 2004). After quality control, normalization (print-tip LOESS; no background correction), and gene specific dye-bias correction (Margaritis et al., 2009) statistical analysis was performed using Limma (Smyth, 2005) for each Cdk8-AA versus the corresponding Parental-AA at the same time point. The reported FC and p-values are an average of the four replicate Cdk8-AA profiles versus the average of four Parental-AA profiles.

### Identifying transient gene expression profiles

Transient expression profiles were identified by correlating the expression profiles of all transcripts in the Cdk8-AA time course to a supervised pre-defined profile. For this the cosine correlation similarity measure was used. A parameter sweep identified the profiles that best fit the data. The parameters represent the time points indicating the start of the peak, the top of the peak (including the width) and when expression returned to Parental-AA. The optimal profiles were identified manually to best fit the data represented in the time course expression profile. The results are robust to small changes in parameters. Transcripts with a correlation better than R=0.9 were considered as part of the cluster.

### Motif discovery and analysis

Putative transcription factor binding motifs were identified for the transient gene expression profiles. The MEME (Multiple Expectation Maximization for Motif Elicitation) (Bailey and Elkan, 1994) algorithm was applied to the sequences encompassing 600 bp upstream of the translation start site of the transient transcripts. Limits for the motifs are set to a 6 basepair minimum and a 20 basepair maximum. Any number of motif repetitions distributed among the sequences was allowed. A third order Markov model was used as background. The most significant motif of each expression profile was submitted to Tomtom for a comparison against a database of known yeast transcription factor binding motifs (Gupta et al., 2007).

### Multiple sequence alignment

The *S. cerevisiae* Gcr2 protein sequences was used in a search against the Refseq protein database of all fungal species using Blastp (Altschul, 1997). Standard settings were used including a word size of 3, BLOSSUM62 as substitution matrix and a gap penalty of 11 for opening and 1 for extending a gap within the alignment. Sequences of the best hits for each species were used to generate a multiple sequence alignment using mafft version 7 (Katoh and Standley, 2013). L-INS-i was used together with the BLOSSUM62 substitution matrix, a gap penalty of 1.53 and offset value of 0.

### Chromatin Immunoprecipitation

ChIP was carried out as previously described (Bakel et al., 2008) with some modifications. In short, 250 mL of mid-log growing yeast cells (OD_595_ = 0.6) were cross-linked with 2% formadehyde for 30 min at 30°C, the reaction was quenched with glycine, and cells were collected by centrifugation. All samples were harvested at the same time and OD. Subsequently, cells were spheroplasted according to the protocol from the Rando lab (Rando, 2010) followed by sonication (Bioruptor, Diagenode: ten cycles, 30 sec on/off, medium setting). 200 μL of chromatin extract was incubated with 10 μL of anti-V5 beads (Sigma) overnight at 4°C. After incubation beads were washed twice in FA lysis buffer (50 mM HEPES KOH at pH 7.5, 150 mM NaCl, 1 mM EDTA, 1% Triton X-100, 0.1% Na-deoxycholate, 0.1% SDS), twice with FA lysis buffer containing 0.5 M NaCl, and twice with 10 mM Tris at pH 8.0, 0.25 mM LiCl, 1 mM EDTA, 0.5% Nonidet P-40 and 0.5% Na-deoxycholate. Cross-linking was reversed overnight by incubating at 65°C in 150 μL 10 mM Tris-HCl (pH 8.0), 1 mM EDTA, 1% SDS. Samples were treated with RNAse and proteinase K, and DNA was recovered for further analysis. The ChIP-chip samples were processed as in (Bakel et al., 2008). For ChIP-qPCR, the fold change of Gcr1 and Gcr2 on the *PYK1* gene promoter was compared to two control regions (HMR-RT and POL1). Primers are listed in Table 3. RT-qPCR was performed using IQ SYBR Green super mix (BioRad) on a 7900HT fast real time PCR machine (Applied Biosystems). All samples were measured as biological triplicates and technical quadruplicates.

**Table 3.**
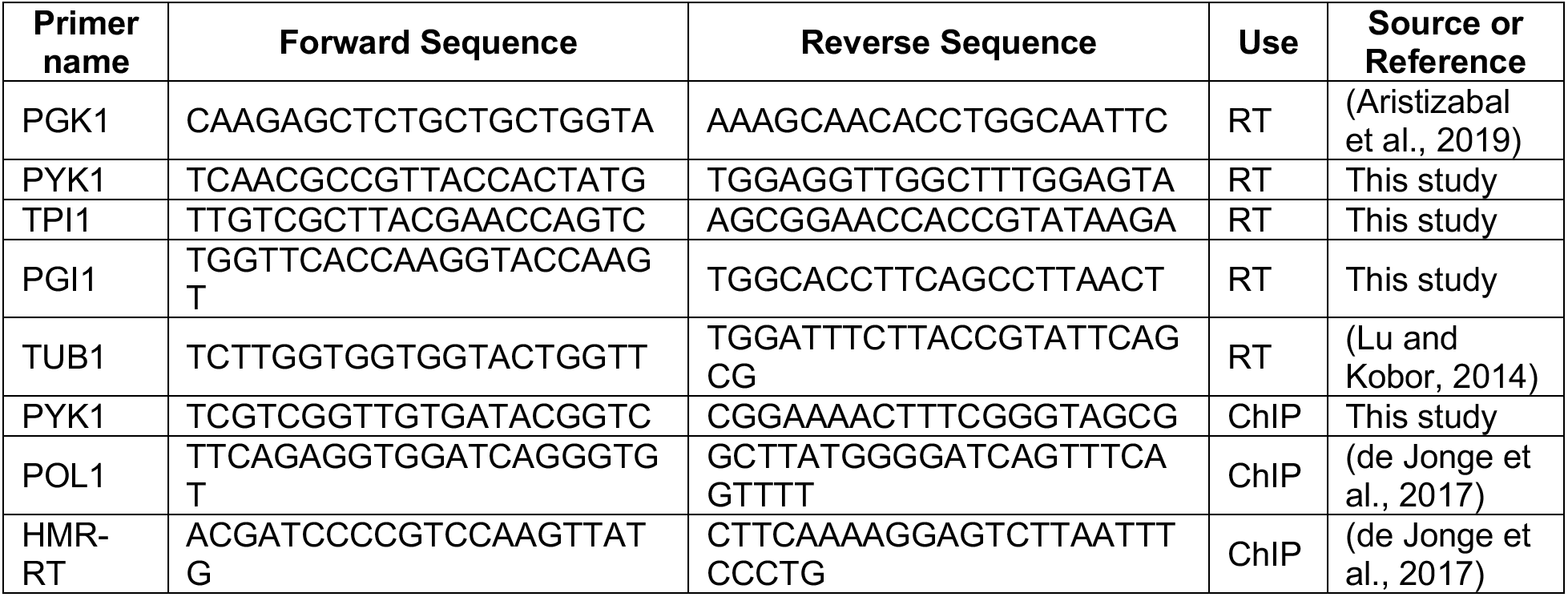
Primers used in this study

### Immunoprecipitation and lambda phosphatase treatment

Overnight cultures were diluted to OD_600_ 0.15 and grown to an OD_600_ of 1.5. For Cdk8 nuclear depletion rapamycin was added to a final concentration of 7.5 μM when cell reached OD_600_ 0.6. Cells were lysed in Lysis buffer (50 mM HEPES (pH 7.5), 150 mM NaCl, 1 mM EDTA, 1% Triton, 0.1% sodium deoxycholate, 0.1% SDS) + Complete protease inhibitor mixture (Roche Applied Science) and 0.1 mM PMSF. Fusion proteins were captured using V5 agarose beads (Sigma # A7345) that were pre-cleared overnight with PBS + 0.1% BSA. Samples were washed twice with Lysis buffer without phosphatase inhibitors and once with 1X NEBuffer for protein metallophosphatases (50 mM HEPES, 100 mM NaCl, 2 mM DTT, 0.01% Brij 35, 1 mM MnCl_2_). Beads were resuspended in 1X NEBuffer for protein metallophosphatases and separated into the indicated treatment groups. 400 units of lambda phosphatase (New England Biolabs, #P0753) was added as indicated, with all samples incubated 15 min at 30°C. Samples were resuspended in Lysis buffer plus 2X SDS buffer and analyzed by SDS-PAGE.

### *In vitro* kinase assays and phosphopeptide analysis

Direct Gcr2 phosphorylation by Cdk8 was tested *in vitro* using FLAG-tagged wild type or kinase deficient (cdk8-D290A) Cdk8 recovered from yeast via immunoprecipitation as described previously (Hirst et al., 1999). Cells were lysed in kinase lysis buffer (KLB) (50 mM Tris, 200 mM NaCl, 5 mM EDTA, 0.1% NP-40, and protease inhibitors), and Cdk8 was immunoprecipitated using anti-FLAG M2 beads. The beads were washed twice with KLB followed by two washes with kinase buffer (KB) (10 mM MgCl_2_, 50 mM Tris [pH 7.5], 1mM DTT, and protease inhibitors). Kinase reactions were performed in 10 μl of KB, 2 pmole of [γ-^32^P]-ATP (Perkin Elmer/ NEN) and 2 μg of recombinant GST, GST-Gcr2, GST-Gcr2(S365A), or GST-CTD substrate protein and incubated for 20 minutes at 30°C. Reactions were resolved by 10% SDS-PAGE and visualized by exposure to Kodak Biomax film.

For analysis of phosphorylated Gcr2 peptides, ^32^P-labled wild type or S356A GST-Gcr2 protein was recovered from SDS-PAGE gels and digested with sequencing grade trypsin (Boehringer Mannheim). Peptides were resolved on cellulose thin layer plates by electrophoresis at pH 1.9 followed by chromatography in butanol/ acetic acid/ H_2_O/ pyridine (75:15:60:50, by volume) in the second dimension. Phosphopeptides were visualized by exposure to Biomax film (Nelson et al., 2003). Phosphoamino acid analysis of tryptic peptides from the *in vitro* kinase reactions was performed as described previously (Sadowski et al., 1991).

### Growth assays

Overnight cultures grown in SC-TRP liquid media were diluted to 0.5 OD_600_, 10-fold serially diluted and spotted onto SC-TRP plates with or without the indicated amounts of hydroxyurea (HU) (Sigma), methyl methanesufonate (MMS) (Sigma), formamide (Sigma), sodium chloride (NaCl) (Bioshop), ethanol, or glycerol. Plates were incubated at the indicated temperatures for 3-5 days.

### RT-qPCR

Overnight cultures were diluted to 0.15 OD_600_ and grown to 0.5 OD_600_ in SC-TRP media. RNA was extracted and purified using the Qiagen RNeasy Mini Kit. cDNA was generated using the Qiagen QuantiTect Reverse Transcription Kit and analyzed using the PerfeCTa SYBR green FastMix (VWR) and a Rotor-Gene 6000 (Qiagen). mRNA levels were quantified from three independent biological replicates using *TUB1* as a control gene. Primer sequences are listed in Table 3.

### Data availability

Complete gene expression and ChIP-chip datasets can be accessed from Gene Expression Omnibus (GEO) with accession numbers GSE166614 and GSE166737 respectively.

## RESULTS

### A conditional depletion strategy to investigate the role of Cdk8 in transcription regulation

Slow growth phenotypes can give rise to altered gene expression profiles that are independent of the underlying genetic alteration (O’Duibhir et al., 2014). Yeast mutants bearing deletion of *CDK8* display slow growth (doubling time of 2.3 hours compared to 1.5 hours in wild type cells) which may confound understanding the role of this kinase in gene regulation. To examine the immediate effects of *CDK8* loss on transcription, we used the anchor-away (AA) system, which causes nuclear proteins to become conditionally sequestered in the cytoplasm by chemical induced multimerization of FKBP12 and FRP fusions in the presence of rapamycin (Haruki et al., 2008). To facilitate comparison of our results with previously generated expression profiles of strains lacking genes encoding mediator subunits or gene-specific transcription factors (Kemmeren et al., 2014; Peppel et al., 2005), we used an anchor away strain generated in the S288C/ BY4742 background (*tor1-1*, *fpr1 RPL13A-FKBP12*)(de Jonge et al., 2017). We confirmed that this strain was insensitive to rapamycin and did not have alterations in global gene expression (Supplementary Figure 1A-D). Furthermore, fusion of a tandem FKBP12-rapamycin-binding (FRB) - GFP tag to the C-terminus of Cdk8 (Cdk8-AA) did not cause growth or gene expression defects under normal growth conditions (Supplementary Figure 1E and F). However, as we expected, exposure of the Cdk8-AA strain to rapamycin caused depletion of Cdk8 from the nucleus within 12 minutes, and at later time points recapitulated the growth defects and flocculation phenotypes of the *cdk8Δ* mutant (Supplementary Figure 1F-H).

### Nuclear depletion of Cdk8 led to altered mRNA levels of specific gene subsets

We examined the effect of nuclear Cdk8 depletion on global mRNA levels over the course of 20 hours using DNA microarrays and observed alterations within 20 minutes of rapamycin treatment (Figure 1, lanes 1 to 23). Importantly, the global mRNA expression profile 20 hours post rapamycin-induced nuclear depletion of Cdk8 was similar to that of a *cdk8Δ* mutant, further validating our approach (Figure 1, lanes 23 and 24, R=0.9). We note that a substantial portion of gene expression alterations identified in this analysis are associated with the slow growth expression profile described previously (compare Figure 1 lane 23 to lane 25, R=0.8) (O’Duibhir et al., 2014). In fact, similarity of expression phenotypes between the Cdk8-AA mutant and the slow growth profile expression patterns gradually increased following addition of rapamycin, reaching a maximum at 20 hours, an effect consistent with the gradual reduction in growth rate observed following Cdk8 nuclear depletion (Supplementary Figure 1F).

**Figure 1:**
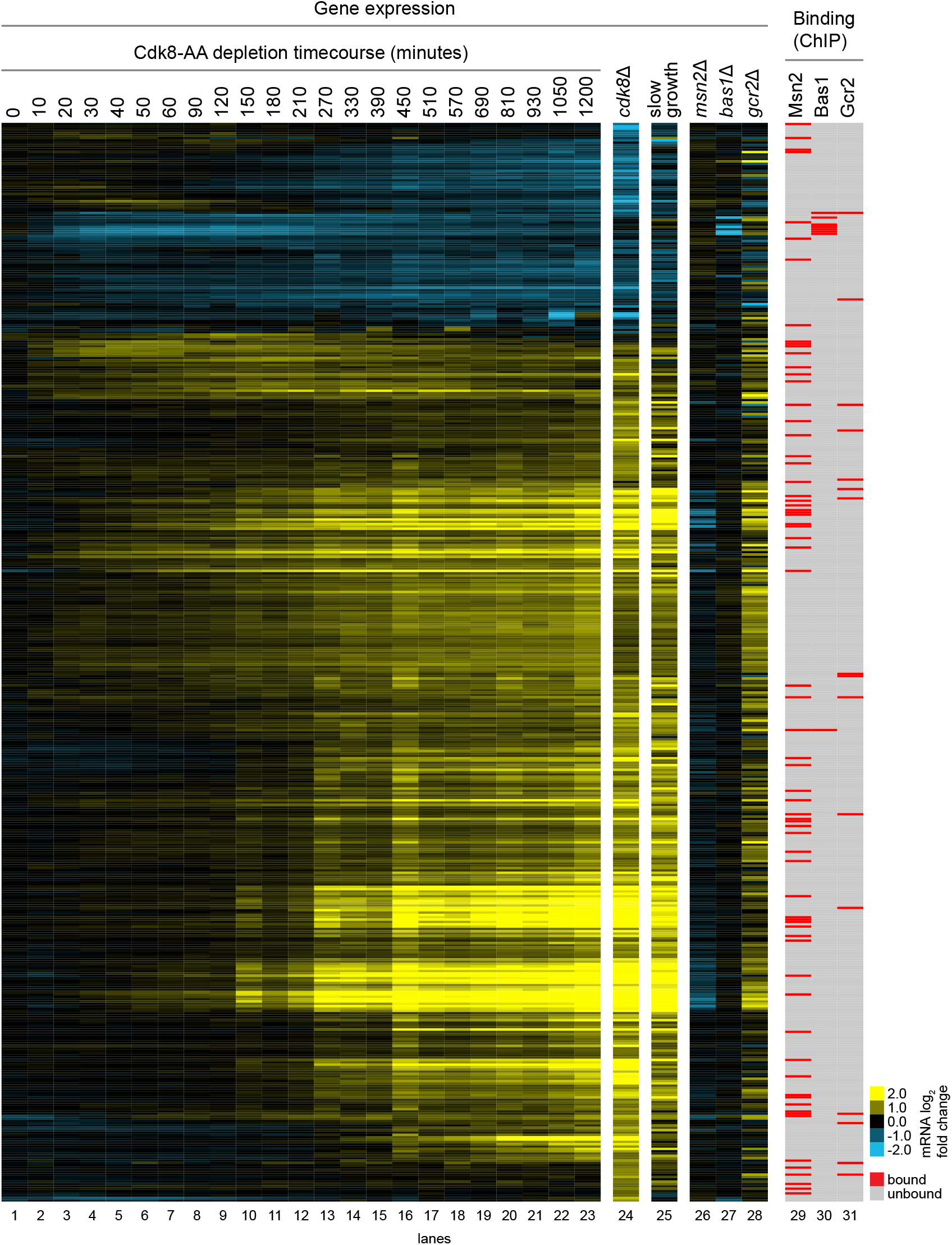
Gene expression alterations upon nuclear depletion of Cdk8. Heat map of gene expression changes and GSTF binding profiles. Genes were cosine clustered based on lanes 1-24 (fold change >1.7 and p<0.05). Lanes 1-23: Time course of gene expression changes following addition of rapamycin to the Cdk8-AA strain. Lane 24: *CDK8* deletion (*cdk8Δ*) profile. Lane 25: Changes associated with slow growth (O’Duibhir et al., 2014). Lanes 26-28: gene expression changes associated with the indicated gene-specific transcription factor deletion strains (Kemmeren et al., 2014). Lanes 29-31: gene-specific transcription factor occupancy (MacIsaac et al., 2006).

To examine direct roles of Cdk8 on gene expression, we focused on alterations that occurred shortly after its depletion. Consistent with previous observations, we identified a cluster of genes whose expression changed upon nuclear depletion of Cdk8 and whose promoter regions were enriched for binding of Msn2 (Hypergeometric test p-value < 0.3 × 10-4), a known transcription factor substrate of Cdk8 (Chi, 2001) (Figure 1, lane 29). These genes have decreased expression in an *msn2Δ* mutant (Hypergeometric test p=4.8 × 10-6), but increased expression upon Cdk8 depletion, an effect in line with a role for Cdk8 in inhibiting Msn2 function (Chi, 2001) (Figure 1). Importantly, nuclear depletion of Cdk8 also produced immediate alterations in the transcription of genes that had not been previously linked to Cdk8 function. For example, we observed a small cluster of transcripts that decreased in abundance early after depletion (20 - 90 minutes), but that returned to untreated levels at later time points (Figure 1, lane 30 overlap with lanes 3-15 p-value < 0.5 × 10-4, hypergeometric test). A focus on these genes revealed that they were enriched for Bas1 binding sites within their promoter regions, and accordingly had decreased mRNA levels in a *bas1Δ* mutant. As such, our conditional depletion approach indicated that Cdk8 may be a positive regulator of this transcription factor, an effect that was missed in earlier *cdk8Δ* mutant gene expression analyses.

### Transcription factor motif analysis highlighted a relationship between Cdk8 and Gcr1/2

To identify additional transcription factors that may be regulated by Cdk8, we focused on genes that showed altered mRNA levels shortly after Cdk8 depletion, and performed transcription factor binding motif analysis on their promoter sequences (Figure 2A). In this analysis we observed enrichment of a motif related to binding sites for Ste12, a known Cdk8 substrate (Nelson et al., 2003) as well as Gcr1 and Gcr2 (Figure 2B), transcription factor partners for the activation of genes encoding glycolysis enzymes (Uemura and Jigami, 1992a). Although Gcr1 and Gcr2 have not previously been linked to Cdk8, work in mammalian cells reported a role for Cdk8 kinase activity in regulating the expression of genes involved in glycolysis, the mechanisms of which remain poorly understood (Galbraith et al., 2017). Analysis of the gene expression pattern of a *gcr2Δ* mutant (Figure 1, *gcr2Δ*, lane 28) revealed that it too was confounded by slow growth (R = 0.7, lane 25 versus lane 28), which may explain why genes whose expression was altered by loss of *gcr2* show no significant enrichment for Gcr2 binding at their promoters (Harbison et al., 2004; MacIsaac et al., 2006). As such, a *bona fide* relationship between Cdk8 and Gcr2 may exist but may have been missed because of confounding effects related to slow growth caused by *cdk8* deletion.

**Figure 2:**
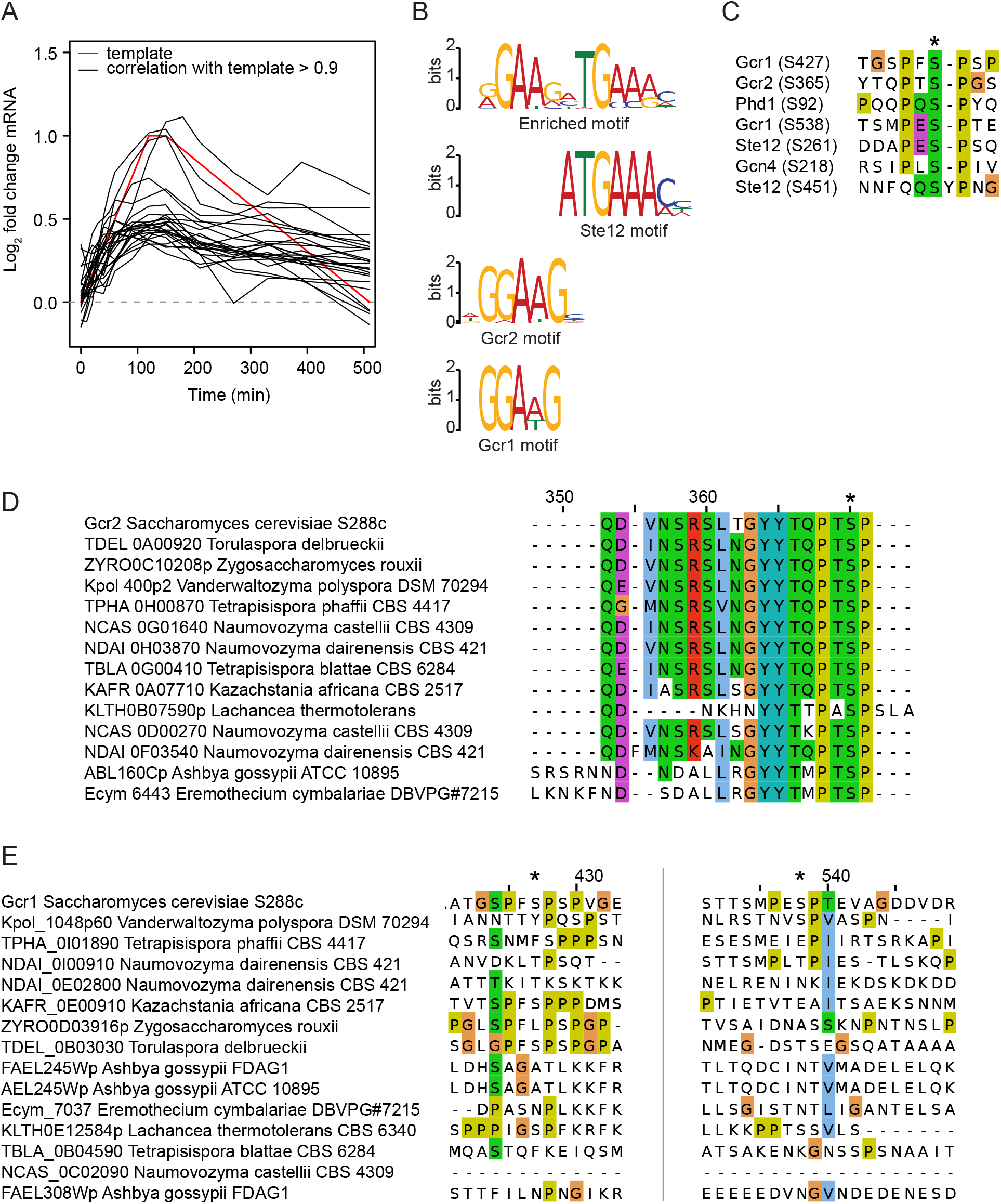
Gcr2 contained a conserved Cdk8 phosphorylation motif. (A) Transcripts whose mRNA level increase shortly after Cdk8 depletion were clustered to an arbitrarily drawn gene expression profile. (B) DNA motif enriched of the gene promoters identified in (A) corresponds to a combined Gcr1/2 and Ste12 motif (from (Gupta et al., 2007)) (C) Potential serine phosphorylation sites (*) on Gcr1 and Gcr2 compared to confirmed sites on Ste12 (Nelson et al., 2003), Phd1 (Raithatha et al., 2011) and Gcn4 (Chi, 2001). (D and E) Multiple sequence alignment displaying homologous Gcr1 and Gcr2 proteins with the highest protein sequence similarity across the indicated fungal species. We found that the serine (*) in Gcr2 that may be targeted for phosphorylation by Cdk8 is highly conserved across species

To determine if Cdk8 may have a direct role in the regulation of Bas1, Gcr1 and/or Gcr2, we examined these proteins for potential Cdk8 phosphorylation sites - serine or threonine residues flanked by a proline 1 or 2 residues toward the C terminus and a proline 2 to 4 residues toward the N terminus (Figure 2C) (Raithatha et al., 2011). No candidate Cdk8 phosphorylation sites were detected in the Bas1 protein sequence, however Gcr1 contained two sites (S427 (PFSP) and S538 (PESP)) and Gcr2 contained one site (S365 (PTSP)) with sequence similarity to previously described Cdk8 phosphorylation sites on Ste12, Phd1, and Gcn4 (Figure 2C). Examining conservation of the residues on Gcr1 and Gcr2 revealed that the potential phosphorylation site on Gcr2 was highly conserved across fungal species (Figure 2D), while the potential sites on Gcr1 were not (Figure 2E). These observations suggest a direct regulatory relationship between Cdk8 and the transcription factors Gcr1 and Gcr2.

### Gcr2 promoter occupancy at glycolysis genes was affected by nuclear depletion of Cdk8

To examine if Cdk8 and Gcr1/Gcr2 functionally interact, we focused our analysis on their target loci. In yeast, glycolysis gene activation is dependent on the transcription factors Rap1, Gcr1 and Gcr2 (Huie et al., 1992; Uemura and Fraenkel, 1990) and consistently, their promoters contain UAS_RPG_ and CT box DNA motifs that are recognized by Rap1 and Gcr1 respectively (Figure 3A). At these sites, Gcr2 does not bind DNA directly, but instead is recruited to glycolysis gene promoters *via* interaction with Gcr1 (Deminoff and Santangelo, 2001; Uemura and Jigami, 1992b). In this complex, Gcr2 may provide transcriptional activation function. Focusing our expression analysis on genes encoding glycolysis enzymes revealed that the loss of *GCR2* resulted in a decrease in their mRNA levels (Figure 3B, lane 26), an effect consistent with previous reports (Uemura and Fraenkel, 1990). Supporting a role for Cdk8 in the regulation of glycolysis gene expression, nuclear depletion of Cdk8 resulted in a progressive increase in the mRNA levels of a subset of glycolysis genes, an effect that was evident 50 min after depletion but that was in part confounded by the slow growth phenotype (Figure 3B, compare lanes 6-23 to lane 25).

**Figure 3:**
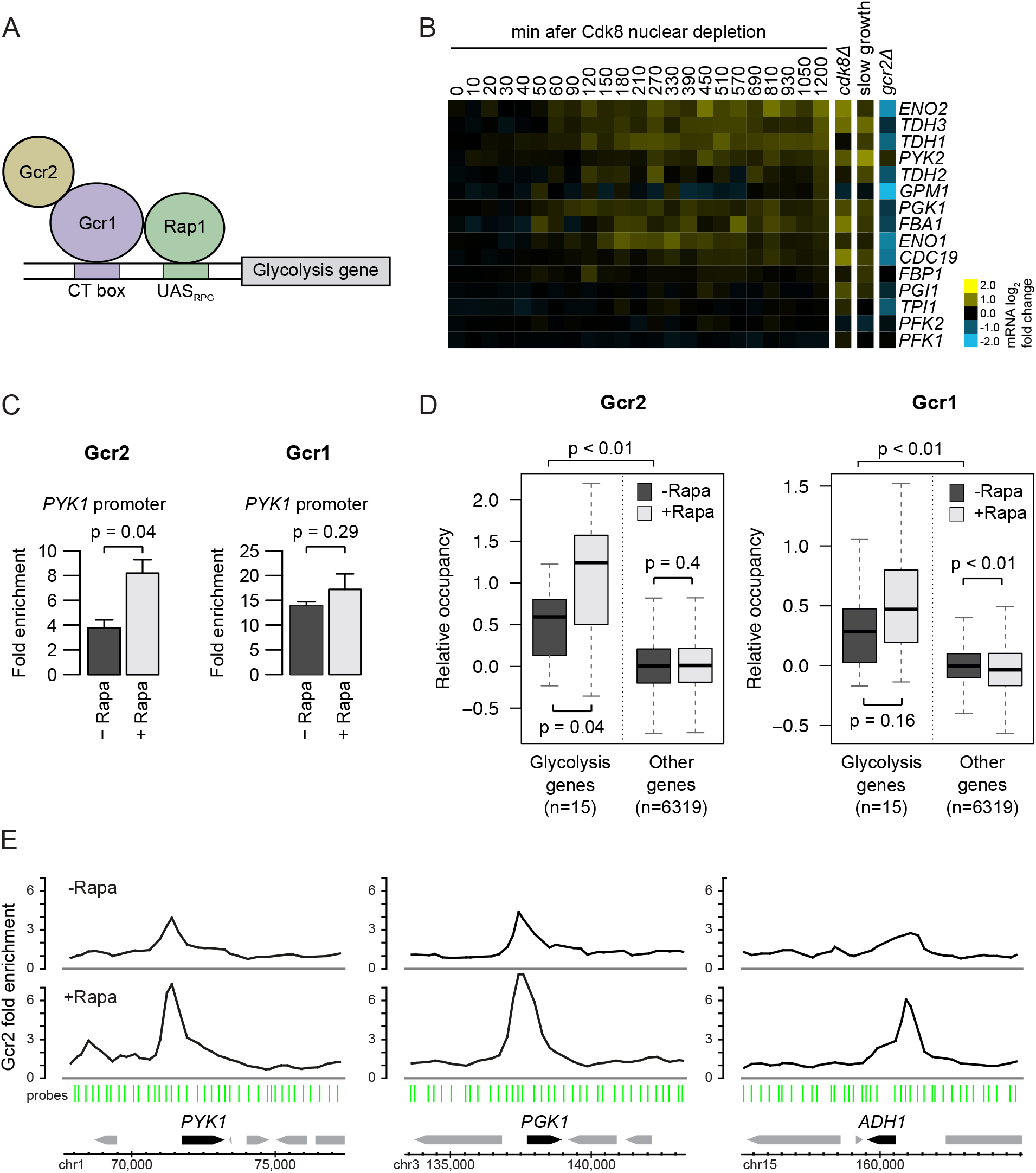
Promoter binding of Gcr2 was dependent on Cdk8 *in vivo*. (A) Schematic of a typical Gcr2-depend gene promoter, highlighting the Rap1 and Gcr1 binding sites. (B) Heat map of gene expression changes upon nuclear depletion of Cdk8 at genes encoding glycolysis enzymes (lanes 1-23). The *cdk8Δ* (lane 24), *grc2Δ* (lane 26) and slow growth (lane 25) gene expression profiles are included for comparison. (C) ChIP-qPCR of Gcr2-V5 (left) and Gcr1-V5 (right) on the *PYK1* promoter before and 60 minutes after Cdk8 nuclear depletion (p values calculated by Student’s t-test). (D) ChIP-chip of Gcr2-V5 (left) and Gcr1-V5 (right) revealed enrichment at genes encoding glycolysis enzymes. Nuclear depletion of Cdk8 for 60 min resulted in a significant increase in Gcr2 occupancy at glycolysis gene promoter, an effect not seen for Gcr1 or at other loci (p values calculated by two-sided Wilcoxon tests). (E) Representative ChIP-chip genomic binding profiles for Gcr2 before and 60 minutes after Cdk8 nuclear depletion.

Finding that the effect of Cdk8 on gene expression extended to Gcr1/ Gcr2-dependent glycolysis genes prompted us to examine if Cdk8 altered the recruitment of Gcr1 and or Gcr2 to glycolysis gene promoters. To this end, we performed chromatin immunoprecipitation and analysis by RT-qPCR. Examination of the *PYK1* gene, which encodes pyruvate kinase, a key enzyme in the glycolysis pathway, revealed that nuclear depletion of Cdk8 resulted in a significant increase in Gcr2 promoter occupancy within 60 minutes (Figure 3C). Expanding this analysis genome wide revealed that genes encoding glycolysis enzymes were enriched for Gcr2 occupancy at their promoters, and that Gcr2 occupancy levels significantly increased upon nuclear depletion of Cdk8 (Figure 3D and E). This effect did not extend to Gcr1, which showed no significant changes in occupancy at the *PYK1* gene promoter or additional genes encoding glycolysis enzymes upon nuclear depletion of Cdk8 (Figure 3C and D). These observations indicate that the interaction of Gcr2 with responsive gene promoters is regulated by Cdk8.

### Gcr2 phosphorylation occurred *in vivo* and was dependent on Cdk8

Cdk8 has well established roles in modulating gene expression *via* phosphorylation of several gene-specific transcription factors (Chi, 2001; Nelson et al., 2003; Raithatha et al., 2011; Rosonina et al., 2012). To determine if Gcr2 is phosphorylated in a Cdk8-dependent manner *in vivo*, Gcr2 was immunoprecipitated from cells with and without nuclear depletion of Cdk8 followed by treatment with or without lambda phosphatase. Treatment of Gcr2 protein recovered from cells without Cdk8 nuclear depletion with lambda phosphatase resulted in the appearance of a faster migrating species compared to untreated samples, indicating the presence of Gcr2 phosphorylation under normal growth conditions (Figure 4A). Consistent with a role for Cdk8 in modulating Gcr2 phosphorylation *in vivo*, we observed a reduction in the slower migrating species upon Cdk8 nuclear depletion, an effect observed within 30 minutes, and which persisted 2 hours after addition of rapamycin (Figure 4A and B). We note that depletion of Cdk8 from the cell nucleus reduced but did not completely eliminate the slower migrating phosphatase-sensitive species of Gcr2, even 2 hours after of rapamycin treatment, findings consistent with multiple residues on Gcr2 phosphorylated *in vivo*, some of which are likely targeted by additional kinases (Albuquerque et al., 2008)

**Figure 4:**
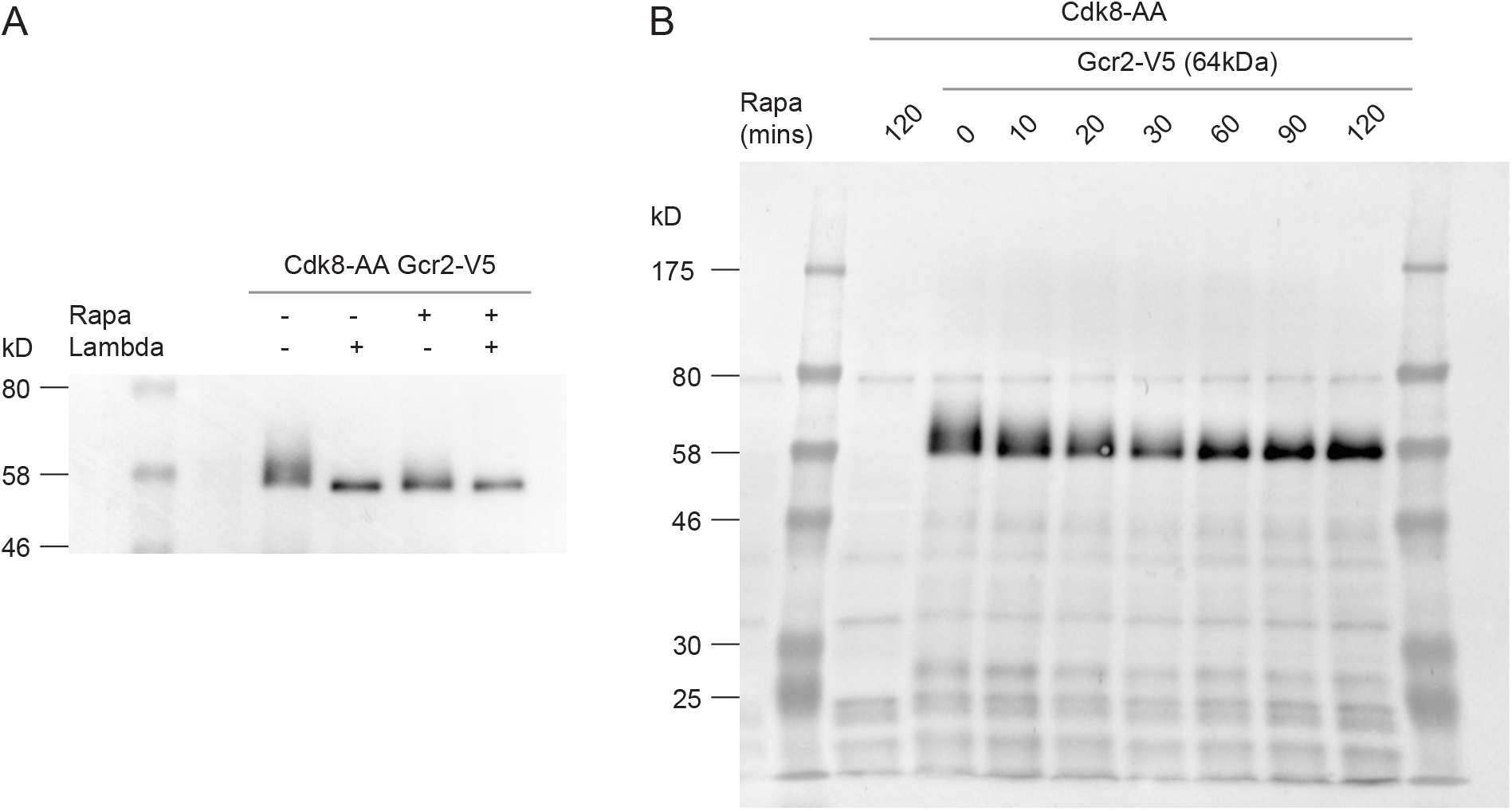
Cdk8 modulated Gcr2 phosphorylation *in vivo*. (A) SDS-PAGE followed by V5 immuno-blot showing lambda phosphatase treatment of V5 immunoprecipitated Gcr2 protein before and 60 min after Cdk8 nuclear depletion. (B) SDS-PAGE followed by V5 immuno-blot of whole cell lysate from a Cdk8-AA/Gcr2-V5 strain during a rapamycin induced Cdk8 nuclear depletion time course.

### Cdk8 directly phosphorylated Gcr2 *in vitro*

Evidence of a role for Cdk8 in Gcr2 phosphorylation *in vivo* (Figure 4A) paired with the presence of a sequence motif on Gcr2 that resembles phosphorylation sites on known Cdk8 substrates (Figure 2C), prompted us to examine if Cdk8 directly phosphorylated Gcr2. For this, we performed *in vitro* kinase reactions using recombinant Gcr2 and Gcr2-S365A mutant proteins (Figure 5A) as substrates for wild type or kinase deficient Cdk8 (D290A) recovered from yeast using a FLAG-epitope tag. In these reactions we found that wild type Cdk8 robustly phosphorylated Gcr2 but less efficiently the Gcr2-S365A mutant (Figure 5B). Importantly, we did not observe phosphorylation of any substrates in reactions with the Cdk8 kinase deficient mutant (D290A).

**Figure 5:**
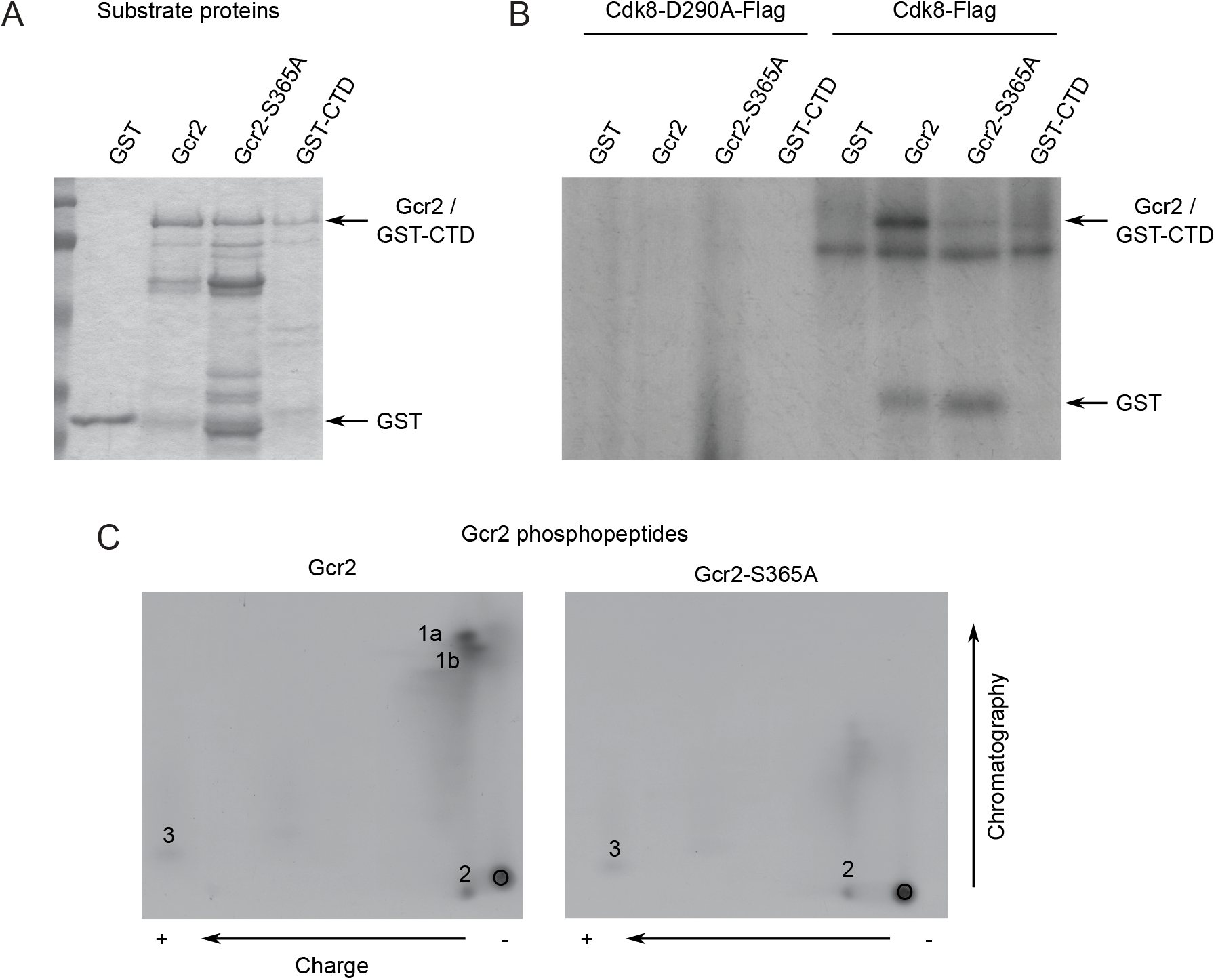
Cdk8 phosphorylated Gcr2 S365 *in vitro*. (A) Gcr2, Gcr2-S365A, GST-CTD and GST were purified from *E. coli* and used for *in vitro* kinase reactions. (B) Flag-tagged wild type (right) or kinase deficient (D290A) (left) Cdk8 was immunoprecipitated from a Σ1278b background and was used for *in vitro* kinase reactions. Gcr2 but not Gcr2-S365A was robustly phosphorylated by wild type but not kinase deficient Cdk8. GST-CTD and GST proteins were used as positive and negative controls respectively. Reactions were analyzed by SDS-PAGE and autoradiography. (C) Phosphopeptide analysis of Gcr2 and Gcr2-S365A proteins phosphorylated by wild type Cdk8. Trypsin digestion of Gcr2 produced four phosphorylated peptides (labelled 1a, 1b, 2, and 3), two of which (1a and 1b) were absent in the Gcr2-S365A mutant.

To determine if Cdk8 specifically targeted S365 of Gcr2, we performed phosphopeptide analysis of Gcr2 and Gcr2-S365A mutant proteins phosphorylated by Cdk8 *in vitro.* We found that Cdk8 kinase-dependent phosphorylation of wild type Gcr2 predominately occurred on a single tryptic peptide (Figure 5C, labelled 1a and 1b) that was not present in reactions with the Gcr2-S365A mutant. Consistently, phosphoamino acid analysis of this phosphopeptide (1a and b) only produced phosphoserine (not shown), effects consistent with S365 being the target site. Nevertheless, we note that phosphorylation of wild type and Gcr2-S365A *in vitro* produced 2 additional phosphopeptides (Figure 5C, labeled 2 and 3), indicating that Cdk8 may phosphorylate additional sites on Gcr2, but with lower efficiency.

### Phosphorylation of Gcr2 at S365 was required for growth on glucose and expression of glycolysis genes

Having established a direct kinase-substrate relationship between Gcr2 and Cdk8 *in vitro*, we examined if Cdk8-dependent phosphorylation of Gcr2 had any significance *in vivo*. To test this, we generated plasmid-based phospho-mutant (*gcr2-S365A*) and phospho-mimic (*gcr2-S365D*) alleles of *GCR2* and introduced them into a *gcr2Δ* mutant background. Strikingly, across several growth conditions the phospho-mutant (S365A) of *GCR2* recapitulated the *gcr2Δ* mutant growth phenotype, an effect not seen with the phospho-mimic allele of *GCR2* (S365D, Figure 6A) and indicative of phosphorylation at S365 being critical for Gcr2 function. To determine if Gcr2 phosphorylation was important for the expression of glycolysis genes, we examined the ability of phospho-mutant and phospho-mimic alleles to sustain expression of representative gene encoding glycolysis enzymes. In line with the growth phenotypes described above, the phospho-mutant allele of *GCR2* recapitulated the *gcr2Δ* mutant gene expression defects, showing decreased mRNA levels for several genes in the glycolysis pathway, effects that were significant for *PYK1* and *PGI1,* with *TPI1* and *PGK1* showing a consistent trend (Figure 6B and C). Again, this effect was not observed with the phospho-mimic version of *GCR2,* indicating that phosphorylation of Gcr2 at S365, a site targeted by Cdk8, was critical for Gcr2’s role as a transcriptional activator of genes encoding enzymes in the glycolysis pathway.

**Figure 6:**
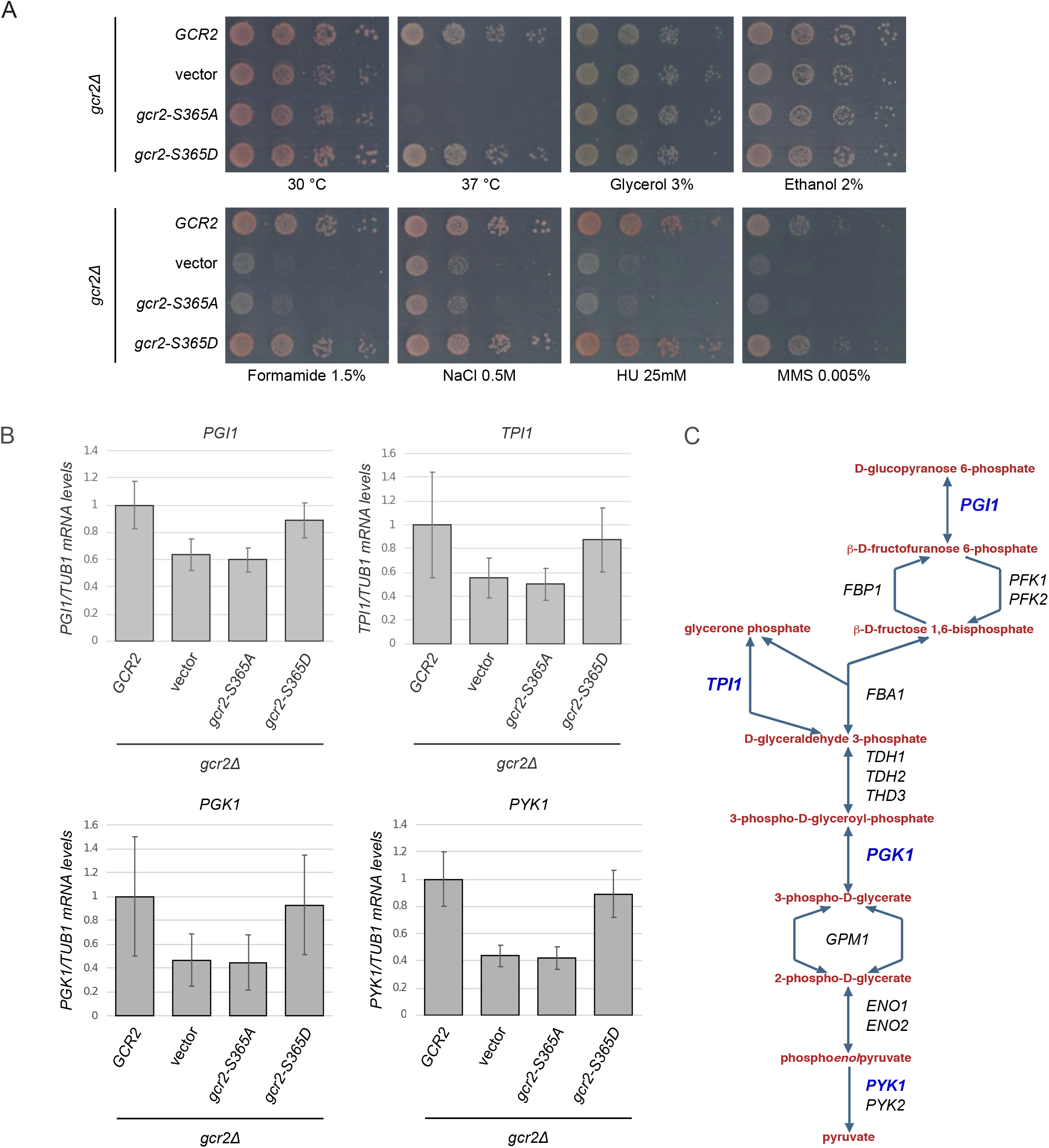
Gcr2-S365 phosphorylation was required for growth and transcription factor function. (A) Loss of Gcr2-S365 phosphorylation recapitulated the *gcr2Δ* mutant growth phenotypes. Cells with the indicated mutations were 10-fold serially diluted, spotted on SC-TRP media and grown for 3-5 days with the indicated carbon source or drug concentrations. (B) RT-qPCR analysis of mRNA levels for representative genes encoding glycolysis enzymes (*PGI1*, *TPI1*, *PGK1 and PYK1*) normalized to *TUB1* mRNA levels (Lu and Kobor, 2014). Error bars represent standard deviation from three independent biological replicates. The *gcr2-S365A* mutant mirrored the *gcr2Δ* mutant gene expression defects at the *PYK1* and *PGK1* genes, underscoring the importance of Gcr2 phosphorylation in gene expression. (C) Schematic of the glycolysis pathway, highlighting the genes tested by RT-qPCR.

## DISCUSSION

The work presented here describes a direct role for Cdk8 in the regulation of genes encoding glycolysis enzymes through phosphorylation of Gcr2, a key transcriptional activator. Using high-resolution gene expression time course profiling following Cdk8 nuclear depletion, we found a connection to transcription factors that were previously not linked to Cdk8: Bas1, Gcr1 and Gcr2. Analysis of the amino acid sequences of these transcription factors identified motifs that mirrored known Cdk8 target sites on both Gcr1 and Gcr2, highlighting their potential as direct Cdk8 substrates. *In vivo* analysis supported the existence of Gcr2 phosphorylation under normal growth conditions and revealed that this modification was dependent on Cdk8 residing in the cell nucleus. A direct kinase substrate relationship between Cdk8 and Gcr2 was supported by *in vitro* kinase reactions, which identified amino acid S365 as a key site on Gcr2 targeted by Cdk8 for phosphorylation. Abolishing phosphorylation of Gcr2 at S365 *via* phospho-mutant alleles revealed that this modification was necessary for normal Gcr2 function, including the activation of genes encoding glycolysis enzymes.

Altogether, our findings uncovered a nuanced mechanism by which Cdk8 modulates Gcr2 and thus the expression of genes encoding glycolysis enzymes. One the one hand, we found that Cdk8 normally functioned to decrease glycolysis gene mRNA levels by limiting Gcr2, and to a lesser extent Gcr1, occupancy at target gene promoters, an effect that persisted 2 hours post nuclear depletion of Cdk8. On the other hand, we found that Cdk8 functioned to activate glycolysis gene expression by targeting Gcr2 S365 for phosphorylation. This discrepancy may reflect the effect of transient vs. stable perturbations on gene expression regulation given that a role of Cdk8 in inhibiting glycolysis gene expression was observed following transient depletion of Cdk8 from the cell nucleus, whereas a role for Cdk8 in stimulating glycolysis gene expression was observed upon stable loss of the main Cdk8 target site on Gcr2, effects that were indistinguishable from the *gcr2* null mutant. Consistently, a role for Cdk8 in activating glycolysis gene expression was reported previously in colon cancer cell lines using constitutive Cdk8 hypomorphic alleles (Galbraith et al., 2017). A dual function for Cdk8 in glycolysis gene regulation is in line with its role in context-specific transcription regulation and may be rooted in its ability to phosphorylate additional sites on Gcr2 as well as other transcription factors, perhaps Gcr1.

In addition to uncovering a new functional relationship between Cdk8 and Gcr2, this work also highlights the utility of conditional depletion strategies paired with dynamic measures of gene expression to uncover mechanisms of transcription regulation that may be missed by traditional approaches using constitutive null mutants. Overall, our work indicates that conditional depletion strategies may be particularly well-suited to separate confounding effects emerging from null mutants, including those associated with slow growth phenotypes which have the potential to mask true associations (O’Duibhir et al., 2014). In this case, depleting Cdk8 from the nucleus using the anchor away system allowed us to focus on the earliest gene expression alterations, which emerged before significant slow growth-associated defects could be detected. Examining rapid gene expression alterations following nuclear depletion of Cdk8, supported a role for Cdk8 in gene-specific transcription regulation. More specifically, within 10-20 minutes post nuclear depletion of Cdk8 we detected increased expression of genes regulated by Msn2, effects consistent with Cdk8 negatively regulating this transcription factor *via* nuclear exclusion (Chi, 2001). Additionally, we detected a set of Bas1-regulated genes whose expression levels were reduced within 20 minutes of Cdk8 nuclear depletion. Given that these genes show high levels of Bas1 occupancy at their promoters and their expression is inhibited by deletion of *BAS1* suggests that Cdk8 may function to stimulate Bas1 activity. Examination of later times points following Cdk8 depletion revealed increasing similarity to the slower growth gene expression profile, which can largely be attributed to downregulation of Gcr2 and upregulation of Msn2 function. Thus, our work suggests that a key role of the Cdk8 kinase is to regulate gene-specific transcriptional regulatory proteins, rather than general components of the transcriptional machinery, for example by phosphorylation of the RNA Polymerase II CTD (Hallberg et al., 2004; Liao et al., 1995; Liu et al., 2004, 2000; Miller et al., 2012; Peppel et al., 2005).

Consistent with the results presented here, previous work in yeast has highlighted both positive and negative roles for Cdk8 on gene-specific transcription regulation *via* phosphorylation of sequence-specific transcription factors. As a negative regulator of transcription, Cdk8 phosphorylates Ste12, Phd1 and Gcn4, targeting them for degradation and leading to the repression of their respective target loci (Nelson et al., 2003; Raithatha et al., 2011; Rosonina et al., 2012). As a positive regulator, Cdk8 phosphorylates Gal4, a modification required for full induction of the *GAL* genes although the exact mechanism remains incompletely understood. Our analysis of cells carrying phospho-mutant alleles of Gcr2 suggests that Cdk8-dependent phosphorylation is necessary for Gcr2’s role as a transcriptional activator, an effect similar to the effect of Cdk8 on Gal4. However, as described above a role for Cdk8 on Gcr2 function is nuanced, also involving transient inhibition of Gcr2 occupancy at target gene promoters.

It is well established that the transcriptional regulation of genes encoding glycolysis enzymes requires the combined activity of several transcription factors: Rap1, Gcr1 and Gcr2. At these genes, Gcr2 associates with target promoter regions *via* direct interaction with Gcr1, which recognizes a CT box motif on the promoter of genes encoding glycolysis enzymes (Huie et al., 1992; Uemura and Fraenkel, 1990). Exactly how Rap1, Gcr1, and Gcr2 function to activate transcription is at present not well understood. Nevertheless, it is clear that transcription of genes encoding glycolysis enzymes requires the recruitment of Gcr1 to CT box motifs and the binding of Gcr2 to Gcr1. Our work indicates that regulation of glycolysis genes also involves decreasing transcription *via* inhibiting Gcr2 promoter occupancy as well as activation of gene expression via Gcr2 phosphorylation at S365 by Cdk8. In the future, it would be of interest to investigate the role of Cdk8 on Gcr1 function and to determine if Gcr2 phosphorylation modulates its ability to interact with Gcr1.

The work described here builds on previous findings in human colon cancer cells lines which showed that Cdk8 kinase activity is required for the activation of glycolysis genes. Although it remains to be determined if the mechanism linking Cdk8 to glycolysis we describe in yeast is conserved in humans, evidence presented here suggests that Cdk8 may function in the regulation of glycolysis across species. As such, our work further emphasizes a key role for Cdk8 in the modulation of metabolism through effects on several pathways including the regulation of genes important for growth on galactose media *via* phosphorylation of Gal4 and low nitrogen conditions via phosphorylation of Gnc4, Ste12 and Phd1. How these activities are coordinated and whether there is any cross-talk between these pathways remains an open question.

## ACKNOWLEDGEMENTS

We thank members of the Holstege, Kobor and Sadowski group for support, discussions and technical assistance.

---

**Supplementary Figure 1:**
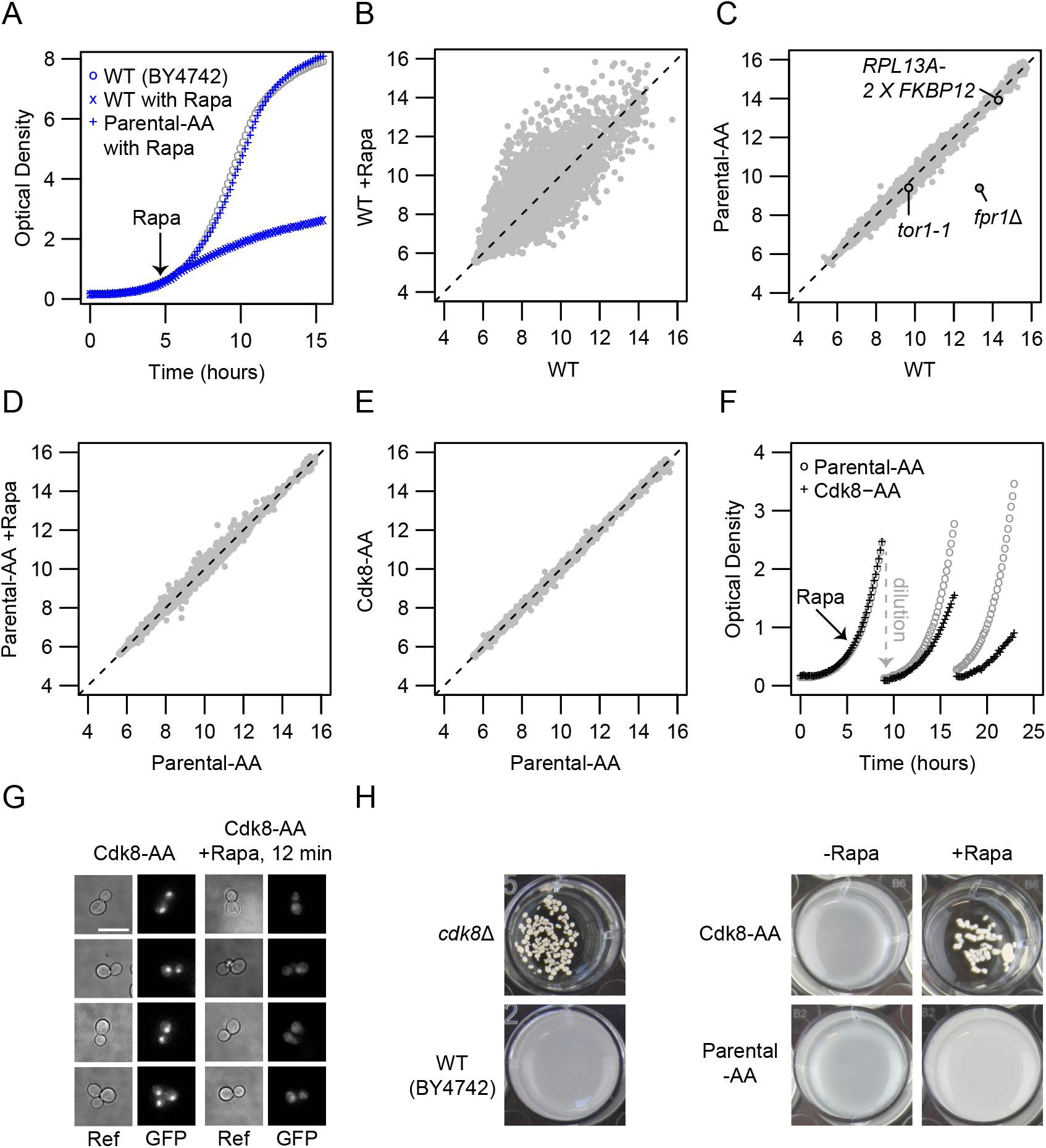
Construction and validation of a conditional Cdk8-AA strain in the BY4742 background. (A) Growth of the Parental-AA strain. (B-E) Scatterplots of gene expression changes of (B) wildtype cells with and without rapamycin treatment; (C) Parental-AA compared to wildtype; (D) Parental-AA treated with and without rapamycin treatment; and (E) Cdk8 tagged with FRB-GFP (Cdk8-AA) compared to Parental-AA. The numbers on the axis refer to the averaged log2 fluorescent dye intensities of the microarray probes representing each gene (dots). (F) Growth of Cdk8-AA strain before and after rapamycin addition. Addition of rapamycin is indicated by the arrow labelled “Rapa”. Cultures were diluted into fresh media with rapamycin twice to avoid nutrient depletion. (G) Fluorescence microscopy of Cdk8-AA (Cdk8-FRB-GFP) before and after rapamycin addition, scale bar 10μM. (H) Exposure of the Cdk8-AA mutant to Rapamycin recapitulates the flocculent phenotype of the *cdk8Δ* mutant.

